# Designed PKC-targeting bryostatin analogs modulate innate immunity and neuroinflammation

**DOI:** 10.1101/2020.09.12.294918

**Authors:** Efrat Abramson, Clayton Hardman, Akira Shimizu, Soonmyung Hwang, Lynda D. Hester, Solomon H. Snyder, Paul A. Wender, Paul M. Kim, Michael D. Kornberg

## Abstract

Neuroinflammation characterizes multiple neurologic diseases, including primary inflammatory conditions such as multiple sclerosis (MS) and classical neurodegenerative diseases. Aberrant activation of the innate immune system contributes to disease progression in these conditions, but drugs that modulate innate immunity, particularly within the central nervous system (CNS), are lacking. The CNS-penetrant natural product bryostatin-1 (bryo-1) attenuates neuroinflammation by targeting innate myeloid cells. Supplies of natural bryo-1 are limited but a recent scalable synthesis has enabled access to it and its analogs (termed bryologs), the latter providing a path to more efficacious, better tolerated, and more accessible agents. Here, we show that multiple synthetically accessible bryologs replicate the anti-inflammatory effects of bryo-1 on innate immune cells *in vitro*, and a lead bryolog attenuates neuroinflammation *in vivo* – actions mechanistically dependent on PKC binding. Our findings identify bryologs as promising drug candidates for targeting innate immunity in neuroinflammation and create a platform for evaluation of synthetic PKC modulators in neuroinflammatory diseases such as MS.

**Graphical Abstract:** 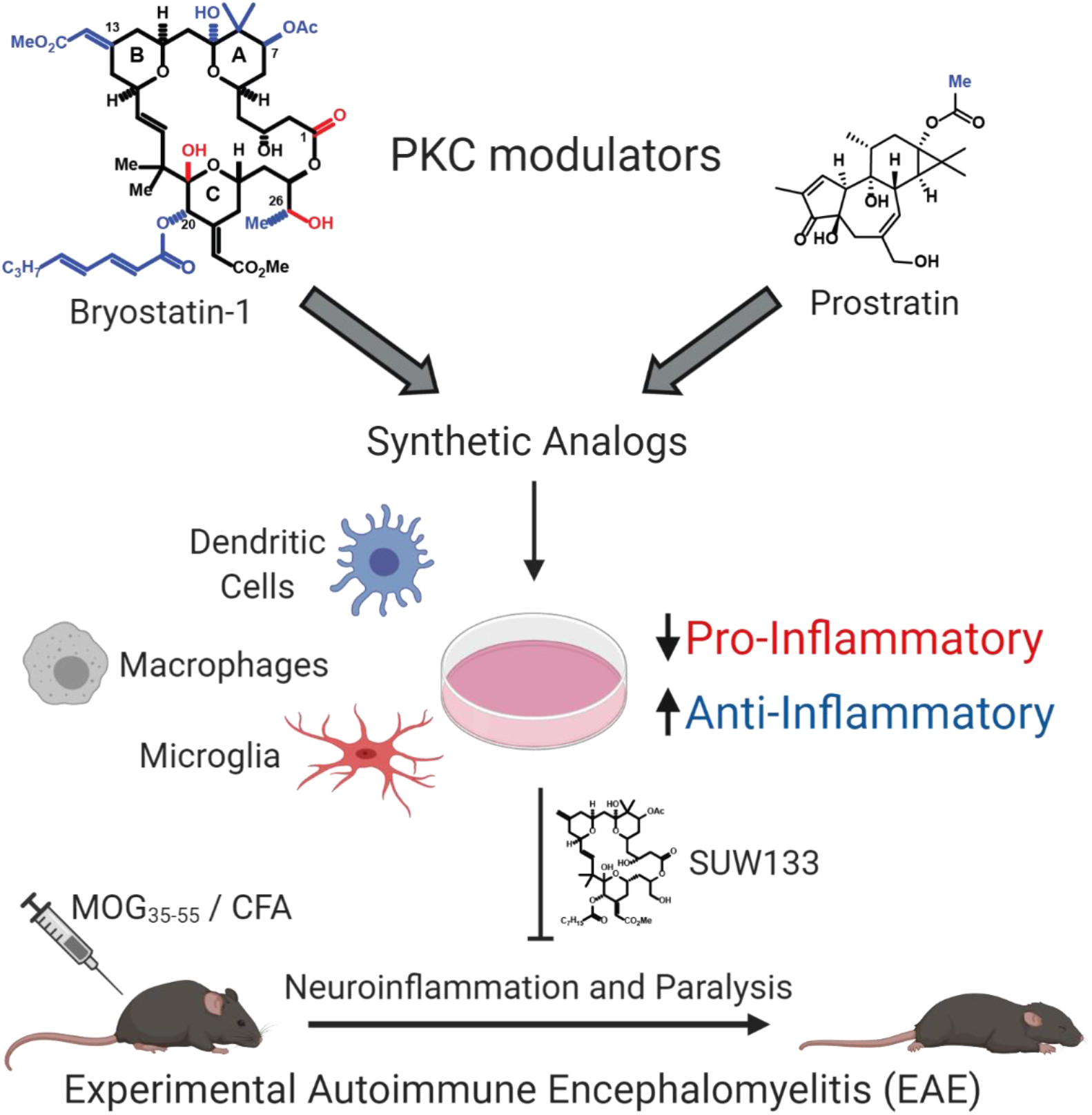

## INTRODUCTION

Neuroinflammation contributes to tissue injury and disability in numerous neurologic diseases, including the prototypic neuroinflammatory condition multiple sclerosis (MS) (Reich, Lucchinetti, and Calabresi, 2018). Although most existing immunomodulatory treatments for MS target the peripheral, adaptive immune system, aberrant activation of innate immune cells plays a prominent role both in the periphery and within the central nervous system (CNS) (Kutzelnigg and Lassmann, 2014; Thompson et al., 2018). In relapsing MS, cells of the innate immune system, such as dendritic cells (DCs), act as antigen presenting cells that drive peripheral lymphocyte activation. In progressive forms of MS, adaptive immunity no longer plays a major role, but compartmentalized activation of innate immune cells (e.g., macrophages and microglia) persists within the CNS and drives neurodegeneration and remyelination failure - the major pathologic substrates of disability (Faissner et al., 2019; Mahad, Trapp, and Lassmann, 2015; Thompson et al., 2018; Voet, Prinz, and van Loo, 2019; Zrzavy et al., 2017). Current therapies fail to modulate innate immunity within the CNS, either because they specifically target lymphocytes or due to poor blood-brain barrier (BBB) permeability, contributing to the lack of successful treatments for progressive MS. Accumulating evidence now implicates a role for microglia in the pathogenesis of many other neurologic diseases, including classic neurodegenerative conditions (Hickman et al., 2018). CNS-penetrant drugs that modulate innate immunity therefore remain a critical unmet need.

Bryostatin-1 (bryo-1) is a naturally occurring, brain-penetrant macrocyclic lactone derived from a marine organism (Pettit et al., 1982; Sun and Alkon, 2006; Trindade-Silva et al., 2010). Pharmacologically, it acts as a potent modulator of protein kinase C (PKC) through its interactions with the diacylglycerol (DAG) binding site of classic (cPKC) and novel (nPKC) isoforms of the enzyme (Ruan and Zhu, 2012). Compared to other PKC modulators that similarly target the DAG binding site, such as plant-derived phorbol esters, bryo-1 produces distinct down-stream effects on PKC function owing to structural elements of the ternary complex formed among bryo-1, PKC, and plasma membrane phospholipids (Ryckbosch, Wender, and Pande, 2017; Wang et al., 1999; Yang et al., 2018). Bryo-1 has anti-tumorigenic properties and has been tested in multiple human clinical trials of solid and hematologic malignancies (Clamp and Jayson, 2002). It is currently under investigation in pre-clinical or clinical studies of Alzheimer’s disease (Farlow et al., 2019; clinicaltrials.gov, NCT03560245), HIV latency reversal (Bullen et al., 2014; Gutierrez et al., 2016; Laird et al., 2015; Sloane et al., 2020), and antigenic stimulation to augment CAR-T immunotherapy (Hardman et al., 2020; Ramakrishna et al., 2019).

We recently reported that natural bryo-1 attenuates neuroinflammation in a mouse model of MS by specifically targeting myeloid cells of the innate immune system, skewing the phenotype of macrophages and DCs from pro-inflammatory to anti-inflammatory/reparative phenotypes (Kornberg et al., 2018). Whether PKC isoforms are the key molecular targets mediating these immunologic actions has remained under investigation, as bryo-1 has also been reported to act directly on toll-like receptor 4 (TLR4) to activate myeloid differentiation primary response 88 (MyD88)-independent signaling through TIR-domain-containing adapter-inducing interferon-β (TRIF) (Ariza et al., 2011). Its targeting of innate immunity, in conjunction with its BBB permeability and established human safety profile, makes bryo-1 an attractive candidate for treatment of neuroinflammatory conditions, including progressive forms of MS that currently lack adequate therapies.

Notwithstanding its promise, it is not clear whether the natural form of bryo-1 is the optimal therapeutic candidate. First, the supply of natural bryo-1 is scarce and variable and isolation from its marine source is difficult, expensive, and an environmental concern (Schaufelberger et al., 1991), creating a barrier to its study and clinical use. Second, as found with most natural compounds (Newman and Cragg, 2020), bryo-1 is not optimized for its molecular/cellular targets and proposed clinical applications, each of which likely depends on distinct PKC isoform(s) and/or other aspects of its pharmacokinetics/pharmacodynamics. Finally, the tolerability window of bryo-1 is narrow, driven by dose-dependent myalgias, creating a need for better tolerated agents with comparable or improved efficacy. As exemplified by taxotere and ivermectin, synthetic analogs of natural products are often clinically superior than their original compounds, with improved efficacy and tolerability (Omura and Crump, 2004; Pazdur et al., 1993). The design, development, and evaluation of synthetic analogs of bryo-1 (termed bryologs) that are synthetically accessible and scalable offer a solution to the supply problem, while enabling exploration of structure-function relationships, that would allow for selection of optimized drug candidates with improved efficacy and limited off-target effects.

We recently reported a scalable synthesis of bryo-1 that addresses the clinical supply problem and provides bryo-1 in quantities that enable the first-ever synthetic access to potentially efficacious close-in precursors and derivatives (Wender et al., 2017). Prior computational, synthesis, and assay work has established the functional importance of the distinct structural elements of bryo-1, identifying pharmacophoric elements critical for PKC binding and regulatory elements that impact ternary interactions with the plasma membrane (**Figure 1A**) (Hardman et al., 2020; Ryckbosch, Wender, and Pande, 2017; Wender et al., 1986 and 1988; Yang et al., 2018). Three hydrogen-bonding functionalities around the C-ring (C1 carbonyl, C19 hemiketal, and C26 hydroxyl) are critical for PKC binding, whereas the A and B rings determine interactions with the plasma membrane and regulate PKC translocation and downstream functions in biological systems. Thus, bryo-1 analogs and other PKC modulators can have similar PKC binding affinities but distinct effects on PKC function. These insights provided the basis for developing a platform of rationally designed bryologs that differ with regard to PKC binding and the kinetics of PKC activation, translocation, and down-regulation within cells, with divergent biological consequences. We recently used this approach to identify a set of bryologs that successfully upregulate cluster of differentiation-22 (CD22) surface expression on leukemia cells (Hardman et al., 2020), an important target for CAR-T immunotherapy (Ramakrishna et al., 2019).

**Figure 1.**
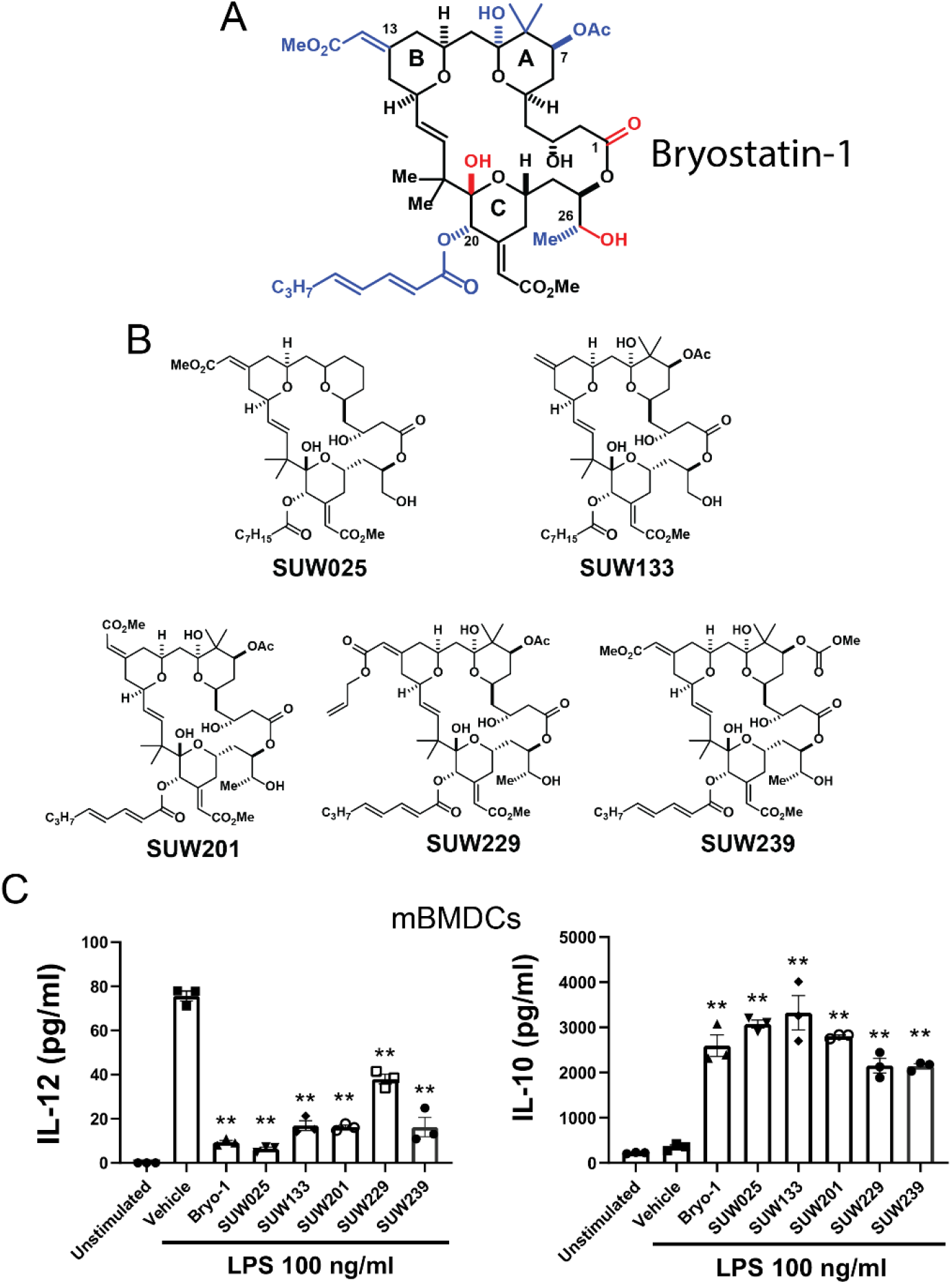
Synthetically Designed Bryologs Replicate the Anti-Inflammatory Effects of Bryo-1 on LPS-Stimulated Dendritic Cells. (A) Structure of bryo-1. Pharmacophoric elements critical for PKC binding are highlighted in red. Nodes for diversification in analog synthesis are highlighted in blue. The C-ring structurally orients the pharmacophoric core, whereas the A- and B-rings regulate PKC interactions with the plasma membrane and downstream functions. (B) Structures of designed bryologs selected for screening in dendritic cells based on PKC binding profiles. (C) Mouse bone-marrow derived dendritic cells (mBMDCs) were left unstimulated or treated overnight with LPS (100 ng/ml) plus vehicle or the indicated bryologs (50 nM). Concentrations of IL-12 (left) and IL-10 (right) were measured from culture media by ELISA assay. Similar to bryo-1, the selected bryologs inhibited IL-12 and augmented IL-10 production. Data represent mean ± SEM of three biological replicates. **p < 0.01 relative to vehicle control, one-way analysis of variance (ANOVA) with Dunnett multiple-comparisons test

Here, we identified a set of synthetically accessible bryologs that replicate the immunologic actions of bryo-1 on innate immune cells. By exploiting structure-function relationships and investigating a structurally unrelated PKC modulator with bryo-1-like effects on PKC in cells, we determined that PKC binding critically mediates the beneficial effects of bryo-1 in cultured myeloid cells and an *in vivo* model of neuroinflammation, experimental autoimmune encephalomyelitis (EAE). We identified a lead bryolog with synthetic advantages and improved *in vivo* tolerability relative to bryo-1 that similarly attenuates disease in the EAE model. These findings clarify the mechanistic target of bryo-1 in neuroinflammation, validate the utility of our chemical platform as both a research tool and therapeutic pipeline, and identify bryologs, and PKC modulators more generally, as promising lead compounds for treating neuroinflammatory conditions such as MS.

## RESULTS

### Identification of synthetically designed PKC modulators that replicate the immunologic actions of bryo-1 in DCs

We previously reported that naturally-derived bryo-1 inhibits production of pro-inflammatory cytokines (such as IL-12) by lipopolysaccharide (LPS)-stimulated murine bone marrow-derived dendritic cells (mBMDCs), while augmenting production of regulatory cytokines such as IL-10 (Kornberg et al., 2018). These actions were associated with the beneficial effects of bryo-1 we observed in autoimmune neuroinflammation. We therefore screened synthetically-derived bryo-1, identical in all respects to the natural product but now more sustainably available, and a set of designed bryologs (with synthetic and/or tolerability advantages) for their ability to replicate the effects of natural bryo-1 in mBMDCs, in order to identify promising lead compounds for further advanced testing.

As noted, prior computational and structure-function studies have determined the functional roles of various elements of the bryo-1 scaffold in PKC binding and regulation (**Figure 1A**) (Ryckbosch, Wender, and Pande, 2017; Wender et al., 1986 and 1988; Yang et al., 2018). The pharmacophoric elements of the C-ring cannot be altered without abrogation of PKC binding, but diversification can be achieved by alteration of regulatory elements in order to improve synthetic accessibility and produce novel drug candidates that can be evaluated for optimized efficacy and tolerability. We therefore examined a number of bryologs with substitutions at C7, C8, C9, C13, C20, and/or C26 (structures shown in **Figure 1B**).These bryologs were chosen on the basis of similar binding affinities to PKC isoforms compared to bryo-1 (**Table 1**), which is necessary but not sufficient to regulate PKC translocation/activation kinetics, along with other potential advantages. SUW025 (synthesis described in DeChristopher et al., 2012) is the simplest and synthetically most accessible bryolog. SUW201, SUW229, and SUW239 (syntheses described in **Supplemental Information** for SUW239 and Hardman et al., 2020 for SUW201 and SUW229) are close-in analogs of bryo-1 selected to determine whether variations in the A- or B-rings would influence immunologic activity as such variations do in other systems. SUW133 (synthesis described in DeChristopher et al., 2012) is a close-in analog chosen based on its established *in vivo* tolerability advantages over bryo-1 (Marsden et al., 2017). This curated set of bryologs had similar effects on LPS-induced cytokine production by mBMDCs compared to bryo-1 (**Figure 1C**).

**Table 1.** PKC Binding Affinities of Bryo-1 Relative to Selected Bryologs and the Prostratin Analog SUW014. Values represent inhibitory constant (Ki, nM) obtained from competitive binding assay with [^3^H]-phorbol-12,13-dibutyrate. Acylation of the C26 hydroxyl group of bryo-1 (SUW275) abrogates binding to all PKC isoforms.

| | PKCα | PKCβI | PKCγ | PKCδ | PKCε | PKCη | PKCθ |
| --- | --- | --- | --- | --- | --- | --- | --- |
| **Bryostatin 1** | 0.81 | 2.2 | 2.2 | 2.1 | 3.0 | 2.2 | 1.5 |
| **SUW025** | 0.91 | 7.49 | 3.45 | 0.94 | 2.24 | 1.64 | 1.59 |
| **SUW133** | 1.8 | 6.3 | 5.6 | 2.0 | 5.8 | 6.4 | 2.0 |
| **SUW201** | 0.54 | 12 | ND | 1.4 | ND | ND | 1.3 |
| **SUW229** | 1.8 | 1 | ND | 1.2 | ND | ND | ND |
| **SUW239** | 2.7 | ND | ND | 1.6 | 0.88 | ND | ND |
| **SUW275** | > 10 μM | > 10 μM | > 10 μM | > 10 μM | > 10 μM | > 10 μM | > 10 μM |
| **SUW014** | 1.4 | 3.1 | 1.9 | 0.5 | 1.7 | 1.9 | 1.6 |

To gain further mechanistic insights, we evaluated a structurally unrelated, designed PKC modulator (SUW014) and a bryolog modified to abrogate binding to PKC (SUW275) (structures shown in **Figures 2A** and **2B**). SUW014 (synthesis described in Beans et al., 2013) is an analog of prostratin, a natural compound structurally related to phorbol esters but with bryostatin-like effects in many systems, including antagonism of phorbol ester-induced tumorigenesis (Gulakowski et al., 1997; Szallasi, Krsmanovic, and Blumberg, 1993). Prostratin is approximately tenfold less potent than bryo-1 as a PKC modulator (Beans et al., 2013; Marsden et al., 2017), while SUW014 has similar PKC binding affinities relative to bryo-1 (**Table 1**). The bryolog SUW275 (synthesis described in Sloane et al., 2020) is “capped” by acylation of the pharmacophorically required C26 hydroxyl group near the C-ring, which as expected abrogates binding to PKC (**Table 1**). Interestingly, the prostratin analog SUW014 similarly replicated bryo-1 actions on mBMDCs, while the PKC non-binding bryolog SUW275 had no effect (**Figure 2C**). These data suggest that PKC binding is critical for the immunologic actions of bryo-1 in DCs, and that structurally unrelated PKC modulators may serve as candidate drugs for targeting innate myeloid cells.

**Figure 2.**
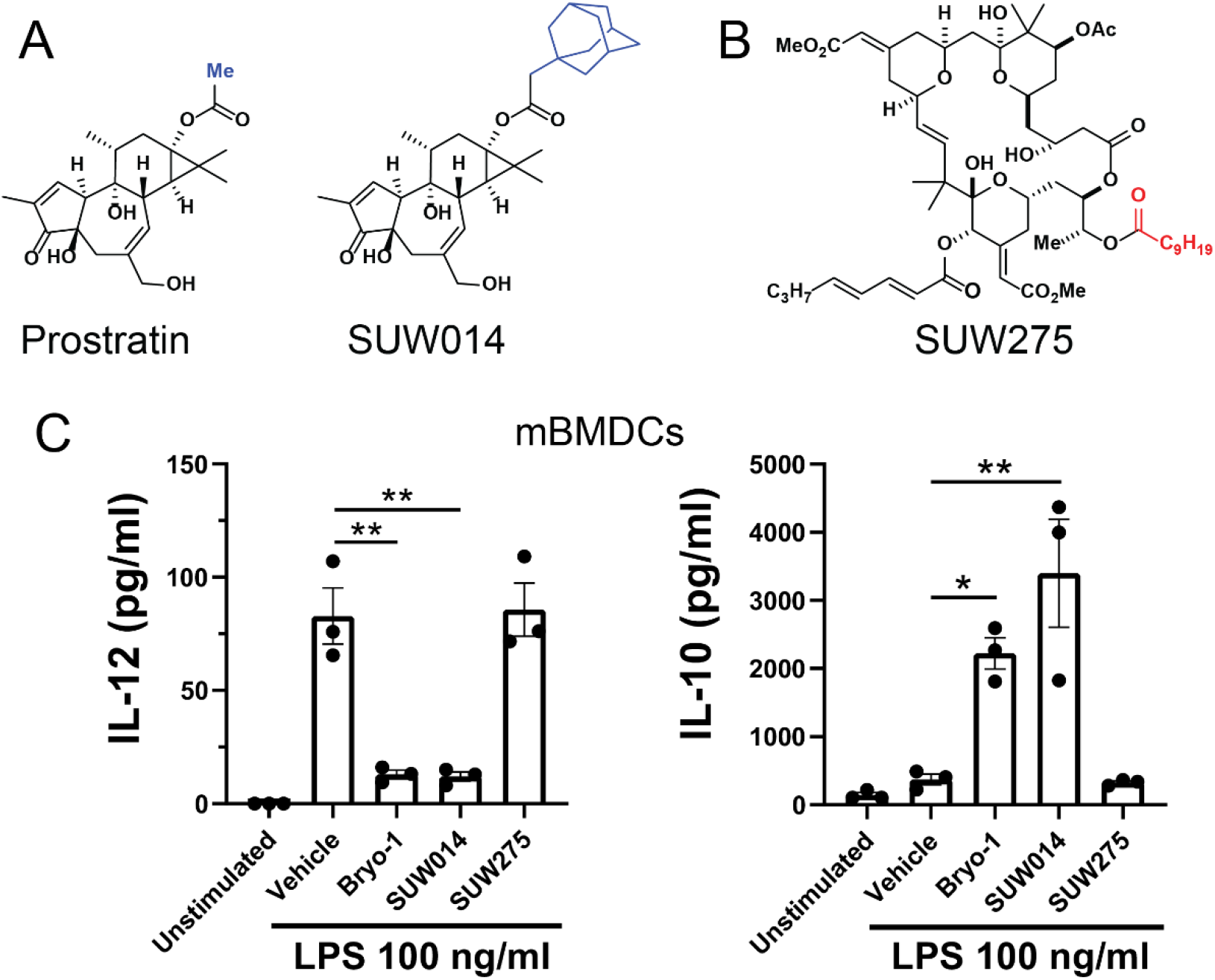
The Immunologic Actions of Bryo-1 in Dendritic Cells Require PKC Binding and are Replicated by a Structurally Unrelated PKC Modulator. (A) Structures of the naturally-derived PKC modulator prostratin and its higher-affinity analog, SUW014. The altered side chain is highlighted in blue. (B) Structure of SUW275, a bryolog designed to abrogate binding to PKC due to acylation of the pharmacophoric C26 hydroxyl group (highlighted in red). (C) mBMDCs were unstimulated or treated overnight with LPS (100 ng/ml) plus vehicle or the indicated compounds (50 nM), followed by quantification of IL-12 (left) and IL-10 (right) secretion. The prostratin analog SUW014, but not the PKC non-binding bryolog SUW275, replicated the effects of bryo-1. Data represent mean ± SEM of three independent experiments performed in triplicate. *p < 0.05, **p < 0.01, one-way ANOVA with Dunnett multiple-comparisons test

### A lead bryolog and prostratin analog replicate bryo-1 effects on macrophages/microglia in a PKC-dependent manner

Macrophages and microglia play critical roles within the CNS during neuroinflammation, determining the delicate balance between tissue injury and repair. We previously reported that bryo-1 inhibits pro-inflammatory activation of macrophages by LPS and augments markers of an “M2-like” reparative phenotype in response to IL-4 (Kornberg et al., 2018). To investigate the potential of synthetic PKC modulators in neuroinflammation, we selected a lead bryolog (SUW133) and evaluated its effects in peripheral macrophages and brain-derived mixed glial cultures, along with the prostratin analog SUW014 and the PKC non-binding bryolog SUW275. SUW133, which is synthetically more accessible than bryo-1 (approximately 25% shorter synthesis), was chosen as a lead compound for further studies based on its substantially greater tolerability window *in vivo* compared to bryo-1 (Marsden et al., 2017 and 2018). Tolerability is the primary limiting factor for bryo-1 in clinical studies. In murine peritoneal macrophages (mPMs), SUW133 and SUW014 inhibited IL-12 production and augmented IL-10 production in response to LPS, similar to bryo-1, while these effects were lost with SUW275 (**Figure 3A**). In mixed glial cultures isolated from mouse brain, SUW133 and SUW014 augmented expression of arginase-1, a marker of “M2-like” phenotype, comparably to bryo-1 after IL-4 stimulation, while SUW275 again had no effect (**Figures 3B** and **3C**).

**Figure 3.**
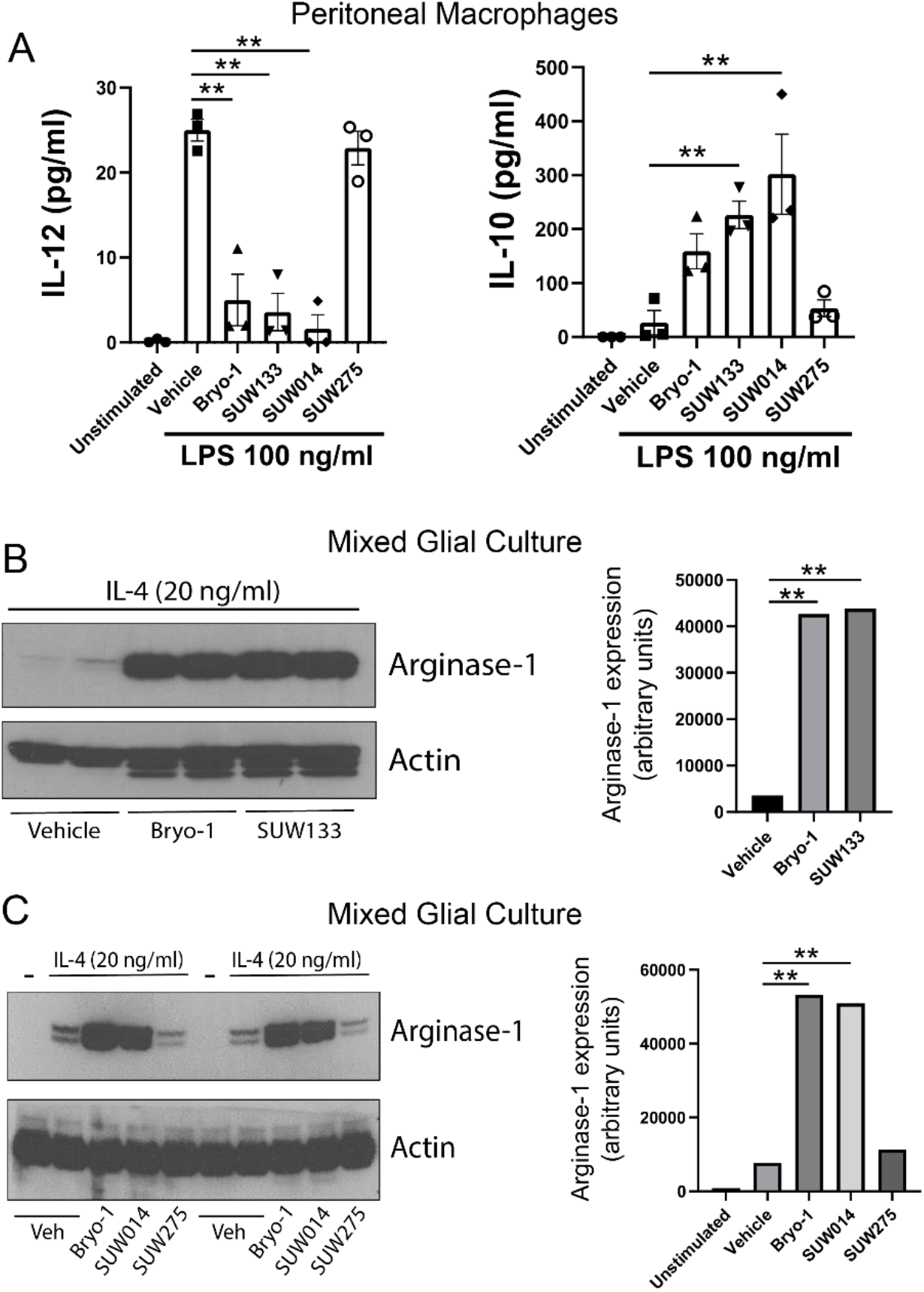
The Lead Bryolog SUW133 and the Prostratin Analog SUW014, but not the PKC Non-Binding Bryolog SUW275, Replicate Bryo-1 Effects on Macrophage and Microglia Phenotype. (A) Peritoneal macrophages were unstimulated or treated overnight with LPS (100 ng/ml) plus vehicle or the indicated drugs (50 nM), followed by quantification of IL-12 (left) and IL-10 (right) secretion. Data represent mean ± SEM of three independent experiments performed in triplicate. (B) Mixed glial cultures were stimulated overnight with IL-4 (20 ng/ml) plus vehicle, bryo-1, or SUW133 (50 nM). Expression of arginase-1, a marker of a regenerative microglial phenotype, was assessed by immunoblot. *Left*, Representative immunoblot. *Right*, Quantification of immunoblots; data represent mean of two independent experiments performed in duplicate. (C) Mixed glial cultures were unstimulated or stimulated overnight with IL-4 (20 ng/ml) plus bryo-1, SUW014, or SUW275 (50 nM), followed by immunoblotting for arginase-1. *Left*, Representative immunoblot. *Right*, Quantification of immunoblots; data represent mean of two independent experiments performed in duplicate. **p < 0.01, one-way ANOVA with Dunnett multiple-comparisons test

Together with the findings from mBMDCs, these data demonstrate that PKC binding is critical for the immunologic actions of bryo-1 in myeloid cells of the innate immune system, both in the periphery and within the CNS, and identify a lead bryolog with synthetic and tolerability advantages that replicates these unique actions. The observation that SUW014, a prostratin analog, produces similar effects suggests that structurally unrelated PKC modulators may serve as candidate drugs for targeting innate myeloid cells.

### Bryo-1 acts independently of TRIF signaling in macrophages and EAE

Our observation that bryo-1 effects on DCs and macrophages/microglia were lost with capping of the PKC-binding pharmacophore (SUW275) and replicated by a structurally unrelated PKC modulator (SUW014) implicates PKC isoforms as the key targets of bryo-1 in innate myeloid cells. Because it has been reported that bryo-1 may act on TLR4 (Ariza et al., 2011) and possibly via the MyD88-independent TRIF signaling pathway, we examined whether the immunologic actions of bryo-1 observed in macrophages and the EAE model of neuroinflammation depend on TRIF. Bryo-1 inhibited LPS-induced IL-12 production equally well in TRIF knockout (KO) compared to wild-type (WT) mPMs (**Figure 4A**), although augmentation of IL-10 production was lost (**Figure 4B**). Active-immunization EAE in C57BL/6 mice is a model of neuroinflammation in which mice are immunized with a peptide derived from myelin oligodendrocyte glycoprotein (MOG35-55) in conjunction with adjuvants, producing robust inflammation in the CNS causing demyelination, axonal injury, and motor deficits. In TRIF KO mice subjected to active-immunization EAE, prophylactic treatment with bryo-1 completely abrogated neurological deficits, similar to the effect seen in WT mice (**Figures 4C** and **4D**). These data suggest that TRIF is not the primary mediator of bryo-1’s immunologic actions in MS models *in vitro* or *in vivo*.

**Figure 4.**
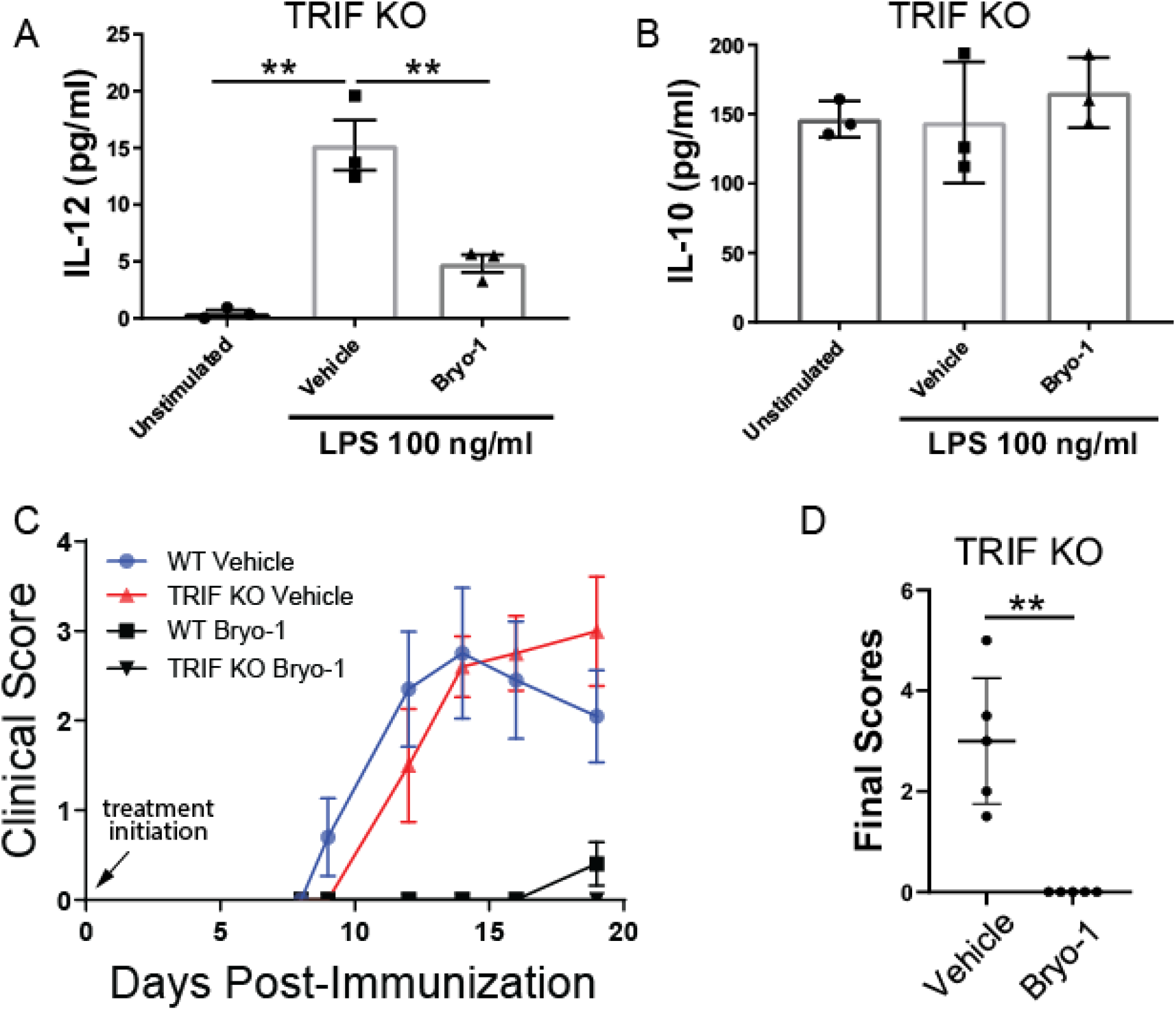
The Anti-Inflammatory Effects of Bryo-1 in Macrophages and EAE are Largely Independent of TRIF. (A and B) Peritoneal macrophages derived from TRIF knockout (KO) mice were unstimulated or stimulated overnight with LPS (100 ng/ml) plus vehicle or bryo-1 (50 nM), followed by quantification of IL-12 (A) and IL-10 (B) secretion. Bryo-1 blocked IL-12 production but failed to augment IL-10 production in TRIF KO mice. Data represent mean ± SEM of three biological replicates. (C) Bryo-1 abrogated EAE independently of TRIF. EAE was induced in wild-type (WT) and TRIF KO mice, and mice were treated with bryo-1 (35 nmol/kg) or vehicle (n = 5 mice per group) via i.p. injection three days per week, beginning on the day of immunization with MOG35-55. Data represent mean ± SEM for each time point. (D) Quantification of final clinical scores of TRIF KO mice treated with either vehicle or bryo-1. Data represent median ± interquartile range of 5 mice per group. **p < 0.01, one-way ANOVA with Tukey multiple-comparisons test in (A), Mann-Whitney *U*-test in (D)

### A lead bryolog attenuates EAE in a PKC-dependent manner

Having established the effectiveness of designed bryologs in immunologic assays of innate myeloid cells *in vitro*, we next evaluated the better-tolerated lead bryolog SUW133 *in vivo* using the EAE model. Active-immunization EAE recapitulates aspects of several neuroinflammatory diseases, including MS, and has proven clinically relevant in MS drug discovery (Robinson et al., 2014). For these experiments, we initiated treatment only after the initial onset of neurological symptoms (therapeutic treatment), corresponding to the early stages of neuroinflammatory injury. Mice were randomized and treatment was initiated once they reached a clinical score of 1.0, corresponding to tail paralysis. As anticipated, SUW133 substantially attenuated neurologic deficits associated with EAE (**Figures 5A** and **5B**). The PKC non-binding bryolog SUW275 had no impact on EAE (**Figures 5C** and **5D**), consistent with our *in vitro* studies and indicating that PKC binding is critical for the immunologic actions of bryo-1 in neuroinflammation *in vivo*.

**Figure 5.**
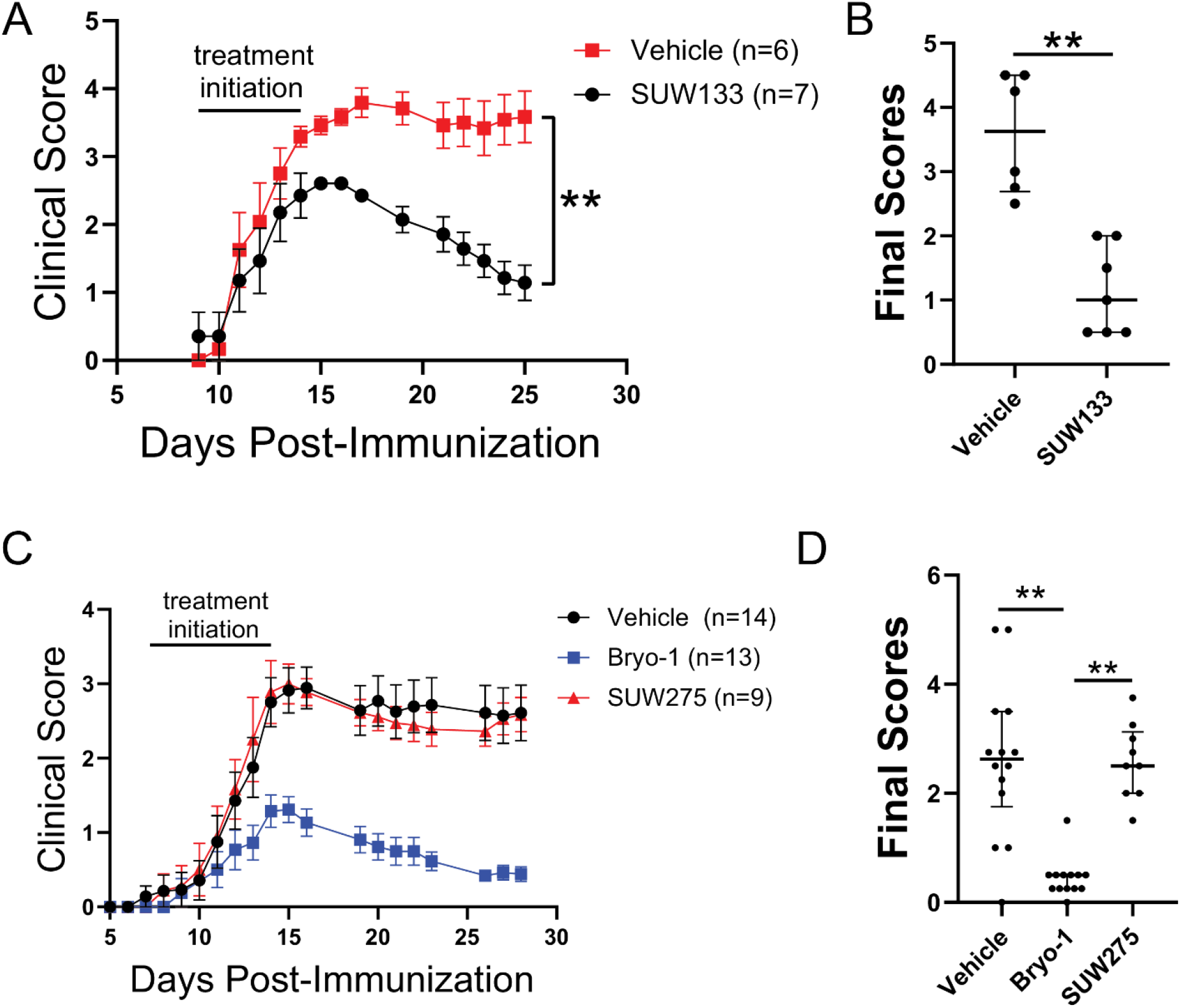
A lead bryolog (SUW133) attenuates EAE in a PKC-dependent manner. (A and B) Mice were subjected to EAE and randomized to treatment with vehicle or the lead bryolog SUW133 (35 nmol/kg) when they reached a clinical score of 1.0 (tail paralysis). Treatment was given three days per week via i.p. injection. Data represent mean ± SEM for each time point in (A). Final clini-cal scores are represented as median ± interquartile range in (B). (C and D) Mice were again subjected to EAE and randomized to treatment groups when they reached a clinical score of 1.0 (tail paralysis). Mice were treated three days per week via i.p. injection with vehicle, bryo-1 (35 nmol/kg), or the PKC non-binding bryolog SUW275 (35 nmol/kg). Data represent mean ± SEM for each time point in (C). Final clinical scores are represented as median ± interquartile range in (D). **p < 0.01, Mann-Whitney *U*-test in (A) and (B), Kruskal-Wallis with Dunn multiple-comparisons test in (D)

## DISCUSSION

Aberrant activation of the innate immune system plays a critical role in a number of autoimmune conditions, including neuroinflammatory conditions such as MS. Yet therapies targeting innate immunity, particularly within the CNS, are lacking. We previously found that bryo-1, a BBB-penetrant natural compound, specifically targets innate immune cells, modulating their phenotypes and attenuating neuroinflammation. But naturally derived bryo-1 has limitations as a drug candidate, including limited and variable supply from natural sources, a narrow tolerability window, and unexplored optimization for clinical applications. Here, we identified a set of rationally designed bryologs that replicate the immunologic actions of bryo-1. These bryologs are synthetically more accessible than bryo-1 and provide a diversified chemical library, allowing screening for candidates with optimized efficacy and tolerability. Immunologic assays in cultured myeloid cells serve as a screening tool to select candidates for further study *in vivo*.

We selected a lead bryolog with a greater *in vivo* tolerability window, SUW133, that showed comparable efficacy compared to bryo-1 in myeloid cell assays and demonstrated its effectiveness in a mouse model of neuroinflammation, providing proof of concept and validating our chemical platform, screening strategy, and assay progression. While decades of research have focused largely on bryo-1 itself, our results showing analogous activity in bryologs and structurally unrelated PKC modulators opens a wide range of research opportunities pertinent to treating neurological disorders.

We found that chemical modifications of bryo-1 designed to abrogate PKC binding also eliminated the immunologic actions of bryo-1 *in vitro* and *in vivo*, identifying PKC as a critical mechanistic target of bryo-1 actions in these systems. This conclusion was further supported by the findings that SUW014, a structurally unrelated PKC modulator, replicated the effects of bryo-1 in *vitro*, and that bryo-1 actions were largely independent of the activation of TLR4’s TRIF pathway. Interestingly, augmentation of IL-10 production by bryo-1 was lost in TRIF KO mPMs. Induction of IL-10 by LPS depends on TRIF signaling (Boonstra et al., 2006; Planes et al., 2016; Teixeira-Coelho et al., 2014), and multiple PKC isoforms modulate the TRIF pathway (McGettrick et al., 2006; Mehta et al., 2012), which likely explains this finding. The observation that bryo-1 maintains efficacy in TRIF KO mice subjected to active-immunization EAE suggests that, in a prophylactic treatment EAE paradigm, effects on IL-10 production may be less important than the inhibition of pro-inflammatory cytokine (e.g., IL-12) production. Myeloid cells transition from a pro-inflammatory to an anti-inflammatory phenotype at the peak of EAE, which is required for the resolution of inflammation (Giles et al., 2018). As such, the effect of bryo-1 on TRIF KO mice may be different when treatment is initiated after the onset of neurologic symptoms, a question we are currently investigating.

Bryo-1 binds to seven distinct isoforms of PKC, and an important future direction is determining which specific isoform(s) mediate bryo-1 actions on the innate immune system and neuroinflammation *in vivo*. Determining the relevant isoforms will allow screening for bryologs and other PKC modulators that are selective for these isoforms, improving tolerability by eliminating off-target effects. Our chemical platform for bryolog synthesis serves as both a research tool and pipeline chemical library in this regard.

In summary, our findings clarify the mechanistic target of bryo-1 in neuroinflammation, identifying PKC as a novel target for modulating innate immunity, and validate synthetic bryologs as promising drug candidates for the treatment of neuroinflammation. Our chemical platform for bryolog development provides the opportunity to identify lead PKC modulating compounds optimized with regard to synthetic accessibility, target selectivity, BBB permeability, and ultimately efficacy and tolerability for the treatment of neuroinflammatory conditions.

## SIGNIFICANCE

Neuroinflammation causes tissue injury and clinical disability in numerous neurologic diseases, including prototypic inflammatory conditions such as multiple sclerosis (MS) and classic neurodegenerative diseases. The innate immune system, both in the periphery and within the central nervous system (CNS), plays a primary role in neuroinflammation, with aberrant responses of CNS-resident macrophages and microglia predominating in progressive forms of MS and neurodegenerative disease. Yet drugs targeting innate immunity, particularly within the CNS, are lacking, presenting a major challenge in the treatment of these conditions. We previously found that bryostatin-1 (bryo-1), a CNS-penetrant natural product known to modulate protein kinase C (PKC), attenuates neuroinflammation through direct effects on innate immune cells, indicating its therapeutic potential. As a naturally-derived compound, bryo-1 is not optimized for clinical applications and has a narrow tolerability window. A need exists for close-in bryo-1 analogs (termed bryologs) that are synthetically accessible with the potential for improved tolerability. Here, by exploiting known structure-function relationships and a chemical platform for generating bryologs, we identified a set of rationally designed, synthetically accessible bryologs that replicate the actions of bryo-1 on innate immune cells in culture. Based on this initial screening, we selected a lead bryolog with known tolerability advantages and demonstrated its efficacy in a mouse model of neuroinflammation. We also show that these immunologic actions are likely mediated by PKC, as they are lost in agents designed to abrogate PKC binding. Our findings implicate PKC as a novel target in neuroinflammation, identify synthetic bryologs as promising lead compounds for modulating innate immunity in neuroinflammatory conditions, and validate our chemical platform as a pipeline for drug discovery.

## ACKNOWLEDGMENTS

This work was supported by grants from the National Institutes of Health (CA31845 to P.A.W., P50 DA044123 to S.H.S.), Conrad N. Hilton Foundation (Marilyn Hilton Bridging Award for Physician Scientists to M.D.K.), and Race to Erase MS (Young Investigator Award to M.D.K.). The authors thank B. Paul, M. Smith, A. Snowman, L. Albacarys, S. McTeer, and P. Calabresi for advice and support.

## AUTHOR CONTRIBUTIONS

Conceptualization, P.A.W., P.M.K, and M.D.K.; Methodology, P.A.W., P.M.K, and M.D.K.; Investigation, E.A., C.H., S.H., L.D.H., P.M.K., and M.D.K.; Formal analysis, E.A., C.H., P.M.K., and M.D.K.; Resources, P.A.W. and S.H.S.; Writing – Original Draft, P.M.K. and M.D.K.; Writing – Review and Editing, E.A., C.H., P.A.W., P.M.K., and M.D.K.; Funding Acquisition, S.H.S., P.A.W., and M.D.K.

## METHODS

| REAGENT or RESOURCE | SOURCE | IDENTIFIER |
| --- | --- | --- |
| **Antibodies** |  |  |
| Mouse monoclonal anti-Arginase-1 | Santa Cruz Biotechnology | Cat# sc-271430 |
| Mouse monoclonal anti-beta Actin, HRP-conjugated | Abcam | Cat# ab20272 |
| **Chemicals, Peptides, and Recombinant Proteins** |  |  |
| Synthetic bryostatin-1 | Wender laboratory | doi: [10.1126/science.aan7969](https://doi.org/10.1126/science.aan7969) |
| SUW025, SUW133 | Wender laboratory | doi:10.1038/nchem.1395 |
| SUW201, SUW229 | Wender laboratory | doi:10.1038/s41467-020-15742-7 |
| SUW275 | Wender laboratory | doi:[10.1073/pnas.1919408117](https://doi.org/10.1073/pnas.1919408117) |
| SUW014 | Wender laboratory | doi:10.1073/pnas.1302634110 |
| SUW239 | Wender laboratory, this paper |  |
| Lipopolysaccharide (LPS) from Escherichia coli O55:B5 | Sigma-Aldrich | Cat# L6529 |
| Recombinant murine IL-4 | PeproTech | Cat# 214-14 |
| MOG<sub>35-55</sub> peptide (amino acid sequence MEVGWYRSPFSRVVHLYRNGK) | Johns Hopkins Synthesis and Sequencing Core Facility | N/A |
| Incomplete Freund's adjuvant | ThermoFisher Scientific | Cat# 77145 |
| Heat-killed Mycobacterium tuberculosis H37 Ra | Difco | Cat# 231141 |
| Pertussis toxin | List Biologicals | Cat# 181 |
| **Critical Commercial Assays** |  |  |
| IL-12 p70 mouse uncoated ELISA kit | Invitrogen | 88-7121-22 |
| IL-10 mouse uncoated ELISA kit | Invitrogen | 88-7105-22 |
| **Experimental Models: Organisms/Strains** |  |  |
| Mouse: Wild-type: C57BL/6J | The Jackson Laboratory | JAX: 000664 |
| Mouse: TRIF knockout: C57BL/6J-*Ticam1*<sup>Lps2</sup>/J | The Jackson Laboratory | JAX: 005037 |
| **Software and Algorithms** |  |  |
| ImageJ | Schneider et al., 2012 | https://imagej.nih.gov/ij/ |
| GraphPad Prism |  |  |
| **Deposited Data** |  |  |
| https://data.mendeley.com/datasets/x33p586t3r/draft?a=0a82eeaf-3da7-4422-bb5a-f3b82631bdcc |  |  |

### EXPERIMENTAL MODEL AND SUBJECT DETAILS

#### Animals

Pregnant female C57BL/6J mice or seven-week-old, female, wild-type C57BL/6J mice (stock # 000664) were purchased from The Jackson Laboratory. Homozygous TRIF knockout mice were also purchased from The Jackson Laboratory (*C57BL/6J-Ticam1^Lps2^*/J, stock # 005037). All mice were housed in a dedicated Johns Hopkins mouse facility. All protocols were approved by the Johns Hopkins Institutional Animal Care and Use Committee.

#### Primary Cell Cultures

##### Murine Bone Marrow-Derived Dendritic Cells (mBMDCs)

mBMDCs were generated as described (Lutz et al., 1999), with minor modifications. Briefly, femurs were removed from 8 – 10 week-old female C57BL/6J mice, cut on both ends, and marrow flushed with PBS. Bone marrow cells were then pelleted by centrifugation (500 x *g* for 8 minutes), re-suspended in red blood cell lysis buffer (Sigma) for one minute, and quenched with excess PBS. After another centrifugation, cells were re-suspended in complete RPMI (cRPMI) media consisting of RPMI-1640 with GlutaMAX supplement (ThermoFisher Scientific) supplemented with 10% fetal bovine serum, penicillin-streptomycin (Gibco), and 50 μM 2-mercaptoethanol (Sigma). On day 0, approximately 2 x 10^6^ cells were then seeded per 100 mm plate in 10 ml of media containing 20 ng/ml of recombinant murine (rm) GM-CSF (Peprotech). On day 3, an additional 10 ml of fresh cRPMI media containing 20 ng/ml rmGM-CSF was added to each plate. On day 6, half the culture supernatant from each plate was removed, centrifuged, and the pelleted cells re-suspended in 10 ml of fresh cRPMI with 20 ng/ml rmGM-CSF and added back to the plates. On day 8, all non-adherent cells (representing the mBMDC fraction) were collected, pelleted by centrifugation, re-suspended in fresh cRPMI media with 20 ng/ml rmGM-CSF, and plated into 24-well dishes. The mBMDCs were then treated overnight with or without LPS (100 ng/ml) plus vehicle or the indicated doses of drug. The following day (day 9), culture media was collected for quantification of cytokine production.

##### Murine Peritoneal Macrophages (mPMs)

Female C57BL/6J mice aged 8 – 12 weeks were injected intraperitoneally with 2 ml each of 3% sterile thioglycollate (BD Biosciences, catalog # 211716) medium. The mice were sacrificed 3 – 5 days later, and cells were harvested by peritoneal lavage with ice-cold RPMI medium containing 2% fetal bovine serum (FBS) and 1 unit/ml heparin. The cells were pelleted by centrifugation at 500 x g for 8 minutes, washed in ice-cold RPMI containing 2% FBS alone, pelleted by centrifugation, and re-suspended in ice-cold Spinner-modification minimum essential medium (SMEM, Sigma, M8167) supplemented with 10% FBS, 2 mM glutamine, and penicillin-streptomycin (SMEM-complete). Cells were then counted, equal numbers were plated into each well of appropriate cell culture dishes, and the cells were incubated at 37°C for 2 - 4 hours to allow macrophage adhesion. Contaminating non-adherent cells were then removed by washing culture dishes five times with ice-cold sterile PBS. Fresh SMEM-complete was then added to the plates, which were incubated overnight at 37°C. The following day, the mPMs were treated overnight with or without LPS (100 ng/ml) plus vehicle or the indicated doses of drugs, followed by quantification of cytokine production from culture media.

##### Mixed Glial Cultures

Mixed glial cultures were prepared from P3-6 C57BL/6 mouse pups. Brains were isolated and placed in EBSS/HBSS in a 60mm culture dish. Meninges were removed and cortices isolated and placed in a 15mL centrifuge tube. A serological pipet was used to remove excess buffer, and 1 mL of 0.05% Trypsin (Gibco Cat #25300) was added per brain. Brains were dissociated in trypsin with a 10 mL serological pipet. Cells were then incubated for 15-20 minutes at 37°C in a water bath. Trypsin was neutralized with equal volume of DMEM/F12 to stop the reaction (Mediatech, Inc. Cat #15-090-CV mixed with 1% antibiotics and 10% FBS). Brains were then homogenized with 6 mL HBSS with a 5 mL serological pipet fixed to a 1 mL sterile pipet tip by pipetting through several times. The cell suspension was filtered twice through a 100 μm cell strainer into a 50 mL conical tube and centrifuged at 1000 rpm for 10 min. Pellet was resuspended in DMEM/F12 media and plated at 250,000 cells/mL on 6-well plates that had been pre-coated with 100 mg/ml PDL (Sigma-Aldrich, cat # 27964-99-4) at 37°C overnight. Media was replaced every three days until confluency at 14 days *in vitro*, at which time the cultures were treated overnight with or without IL-4 (20 ng/ml) plus vehicle or the indicated drugs, followed by evaluation of arginase-1 expression by immunoblotting.

### METHOD DETAILS

#### Synthesis and Characterization of Analogs

All reactions were conducted in oven- or flame-dried glassware under a nitrogen or argon atmosphere unless otherwise noted. Reactions were concentrated under reduced pressure with a rotary evaporator unless otherwise noted. Commercial reagents were used as received or purified using the methods indicated herein. Dichloromethane, diethyl ether, dimethylformamide, pentane, tetrahydrofuran, and toluene were passed through an alumina-drying column (Solv-Tek Inc.) using nitrogen pressure; ethyl acetate, hexanes, and petroleum ether were obtained from Fisher Scientific. Analytical thin-layer chromatography (TLC) was carried out on 250 μm silica gel 60G plates with fluorescent indicator F254 (EMD Millipore). Plates were visualized with UV light and treated with *p*-anisaldehyde, ceric ammonium molybdate, or potassium permanganate stain with gentle heating. Flash column chromatography was performed using silica gel (230-400 mesh, grade 60, particle size 40 to 63 μm) purchased from Fisher Scientific. pH 7 buffered silica gel was prepared by adding 10% weight pH 7 phosphate buffer to silica and rotating for ~12 hrs. NMR spectra were acquired on a Varian INOVA 600, Varian INOVA 500, or Varian 400 magnetic resonance spectrometer. ^1^H chemical shifts are reported relative to the residual solvent peak (CHCl_3_ = 7.26 ppm, C_6_H_6_ = 7.16 ppm) as follows: chemical shift (δ), multiplicity (app = apparent, b = broad, s = singlet, d = doublet, t = triplet, q = quartet, m = multiplet, or combinations thereof), coupling constant(s) in Hz, integration. ^13^C chemical shifts are reported relative to the residual solvent peak (CHCl_3_ = 77.16 ppm, C_6_H_6_ = 128.06 ppm). Infrared spectra were acquired on a Nicolet iS 5 FT-IR Spectrometer (ThermoFisher). Optical rotations were acquired on a P-2000 Digital Polarimeter (Jasco). High-resolution mass spectra (HRMS) were acquired at the Vincent Coates Foundation Mass Spectrometry Laboratory at Stanford.

Synthesis and characterization of SUW239 are shown in **Supplemental Information**. Synthesis and characterization of the other analogs were previously reported (DeChristopher et al., 2012 for SUW025 and SUW133; Hardman et al., 2020 for SUW201 and SUW229; Beans et al., 2013 for SUW014; Sloane et al., 2020 for SUW275).

#### PKC Binding and PKCδ-GFP Translocation Assays

These assays were performed as described in detail in Supplemental Information from Sloane et al., 2020.

#### Drug Preparation and Delivery

All drugs were initially prepared as 10 mM stock solutions in DMSO. For treatment of cultured cells, drugs were diluted in 100% ethanol to a final concentration of 50 μM to create working solutions, which were stored at 4°C for up to 3 months. Cultured cells were then treated with drug at 1:1,000 dilution to a final concentration of 50 nM, or with equal volume of 100% ethanol as vehicle control. For EAE experiments, stock solutions were diluted to 7 nmol/ml in 20% ethanol in PBS to create working solutions, which were stored at −20°C until use. Mice were treated with working solutions via intraperitoneal injection, with 20% ethanol in PBS as vehicle control.

#### Quantification of Cytokine Production by Enzyme-Linked Immunosorbent Assay (ELISA)

After overnight treatment of mBMDCs or mPMs with LPS 100 ng/ml +/- vehicle or drug, culture supernatants were collected and cytokine production was assayed using ELISA kits for IL-12 p70 (catalog # 88-7121-22) and IL-10 (catalog # 88-7105-22) purchased from eBioscience, according to manufacturer’s instructions. Plates were read at 450 nm on a table-top plate reader.

#### Cell Lysate Preparation and Immunoblot Analysis from Mixed Glial Cultures

Mixed glial cultures were washed with PBS, and lysates were prepared by incubation in ice-cold RIPA buffer (50 mM Tris pH 7.4, 150 mM NaCL, 1% triton, 0.5% sodium deoxycholate, 0.1% SDS, 1 mM EDTA) supplemented with protease inhibitors. Lysates were cleared by centrifugation, and protein concentration was measured by Bradford assay. Lysates were mixed with SDS sample buffer, boiled, and resolved by SDS-PAGE. Bands were transferred to PVDF Immobilon P membranes (Millipore) using a wet transfer, blocked in TBS-T containing 5% milk, and probed overnight at 4°C with primary antibody against arginase -1. Membranes were then incubated for 1 hour with HRP-conjugated secondary antibody (Jackson ImmunoResearch). Immunoblots were visualized using the SuperSignal West ECL system (Thermo Scientific) followed by film exposure. Blots were then stripped using Restore Western blot stripping buffer (Thermo Scientific, catalog # 21059), blocked again as above, and probed overnight at 4°C with primary antibody against actin, prior to visualization as above. Quantification was performed using ImageJ software.

#### Induction and Scoring of Experimental Autoimmune Encephalomyelitis (EAE)

Active EAE was induced in 8–12 week-old female C57BL/6J mice that had been allowed to acclimatize to the animal facility for at least one week. MOG35-55 peptide dissolved in PBS at a concentration of 2 mg/ml was mixed 1:1 with complete Freund’s adjuvant to make an emulsion. On day 0, mice were immunized by injecting 50 μl of the emulsion subcutaneously into each of two sites on the lateral abdomen. In addition, on day 0 and again on day 2, mice were injected i.p. with 250 ng of pertussis toxin dissolved in PBS. Mice were weighed and scored beginning on day 7 post–immunization. Scoring was performed by a blinded observer according to the following scale: 0 = no clinical deficit; 0.5 = partial loss of tail tone; 1.0 = complete tail paralysis or both partial loss of tail tone plus awkward gait; 1.5 = complete tail paralysis and awkward gait; 2.0 = tail paralysis with hind limb weakness evidenced by foot dropping between bars of cage lid while walking; 2.5 = hind limb paralysis with little to no weight-bearing on hind limbs (dragging) but with some movement possible in legs; 3.0 = complete hind limb paralysis with no movement in lower limbs; 3.5 = hind limb paralysis with some weakness in forelimbs; 4.0 = complete tetraplegia but with some movement of head; 4.5 = moribund; 5.0 = dead.

#### Drug Treatment of EAE Mice

Mice were treated 3 days per week via intraperitoneal injection with drug or vehicle control (20% ethanol in PBS). For the experiment comparing bryo-1 effects in wild-type versus TRIF knockout mice (Figures 4C and 4D), treatment began on the day of EAE immunization (prophylactic treatment). For experiments with bryologs (Figure 5), mice were randomized to treatment groups once they reached a clinical score of 1.0, corresponding to tail paralysis (therapeutic treatment). Mice were randomized by an observer blinded to subsequent treatment groups.

### QUANTIFICATION AND STATISTICAL ANALYSIS

All statistical analyses were performed using GraphPad Prism software. Quantification of immunoblots was performed with ImageJ software. Details of statistical analyses for each experiment can be found in the figures and figure legends.

