## Supplemental Information for "Designed PKC-targeting bryostatin analogs modulate innate immunity and neuroinflammation"

### Supplemental Information (SI)

#### SYNTHESIS AND CHARACTERIZATION OF SUW239

#### *Synthesis of C7-modified bryostatin analogs: C26 Alloc protection of bryostatin 1*

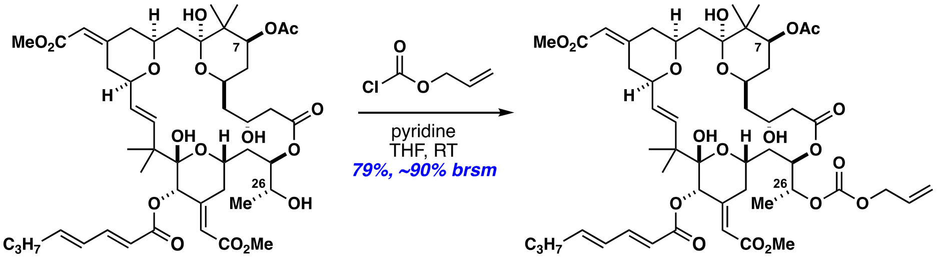


B1 1

To a flame dried flask containing bryostatin 1 (58 mg, 0.064 mmol, 1 equiv.) and a magnetic stir bar was added THF (1 mL, ~0.064M) and pyridine (0.045 mL, 0.55 mmol, ~9 equiv.) and cooled in a water/ice bath. To this allyl chloroformate (0.026 mL, 0.249 mmol, ~4 equiv.) was added and the reaction mixture was allowed to warm to room temperature and stirred for 18 hours. At this point the solution was slightly cloudy and TLC of the reaction mixture showed incomplete conversion. To this an additional portion of allyl chloroformate (0.026 mL, 0.249 mmol, ~4 equiv.) was added and the solution was stirred at room temperature for an additional 4 hours. The solution was quenched into 40 mL of DI water and extracted with EtOAc (4 portions of 40 mL) and the resulting organic layer was washed with 4:1 Brine:1N HCl (20 mL). The acidic brine wash was back-extracted with 40 mL EtOAc. The combined organic layers were dried over Na2SO4, filtered, and concentrated to a crude film. Purification was accomplished by silica gel flash column chromatography (20%-40%-60%-100% EtOAc/Hex) affording product 3.14 (50.2 mg, 79% yield) and recovered starting material bryostatin 1 (12 mg, 20% of starting material). Compound purity was established by TLC (one spot) analysis.

***Characterization data for C26-Alloc protected bryostatin 1***

**TLC $R_f$** = 0.83 (60% EtOAc/hexanes, purple spot in p-anisaldehyde)

**$[\alpha]^{22.3}_{D}$** = 41.41° ± 0.53° (c = 0.155 wt/vol%, CH2Cl2)

**IR** (thin film) 3466, 3319, 2950, 2930, 2850, 1742, 1717, 1652, 1645, 1616, 1436, 1367, 1245, 1162, 1153, 1099, 1030, 1003, 859, 812 cm-1

**$^{1}$H NMR** (500 MHz, Chloroform-d) δ 7.30 – 7.20 (m, 1H), 6.21 – 6.12 (m, 2H), 5.99 (d, J = 2.0 Hz, 1H), 5.99 – 5.88 (m, 1H), 5.80 (d, J = 2.0 Hz, 1H), 5.77 (d, J = 2.4 Hz, 1H), 5.66 (t, J = 1.8 Hz, 1H), 5.42 – 5.25 (m, 4H), 5.18 (bs, 2H), 5.14 (dd, J = 11.8, 4.8 Hz, 1H), 4.94 – 4.84 (m, 1H), 4.64 (dd, J = 5.9, 1.4 Hz, 2H), 4.30 – 4.05 (m, 4H), 3.99 (tt, J = 11.2, 2.8 Hz, 1H), 3.82 – 3.74 (m, 1H), 3.69 (s, 3H), 3.65 (s, 3H), 3.70 – 3.59 (m, 2H), 2.58 (s, 1H), 2.49 – 2.37 (m, 2H), 2.24 – 2.05 (m, 6H), 2.04 (s, 3H), 2.03 – 1.86 (m, 3H), 1.82 (td, J = 14.1, 11.3, 2.9 Hz, 1H), 1.75 (dt, J = 12.0, 4.3, 2.3
Hz, 1H), 1.66 (d, J = 15.1 Hz, 1H), 1.58 (dt, J = 14.4, 2.8 Hz, 1H), 1.51 – 1.39 (m, 3H), 1.29 (d, J = 6.5 Hz, 3H), 1.13 (s, 3H), 0.99 (s, 6H), 0.94 (s, 3H), 0.91 (t, J = 7.4 Hz, 3H)

**$^{13}$C NMR** (125 MHz, CDCl3) δ 171.0, 170.9, 167.1, 166.8, 165.7, 156.3, 154.6, 152.0, 146.5, 145.7, 139.4, 131.6, 129.5, 128.5, 119.7, 119.3, 118.7, 114.7, 101.8, 99.2, 79.1, 75.3, 74.1, 72.8, 71.5, 70.0, 68.8, 68.7, 65.9, 64.7, 51.2 (2 Carbons), 45.0, 44.2, 42.3, 42.1, 41.1, 40.1, 36.4, 35.6, 35.2, 33.5, 31.3, 24.7, 22.0, 21.3, 21.1, 19.9, 16.9, 16.3, 13.8

**HRMS:** calculated for C51H72NaO19 [M+Na]+: 1011.4560; found 1011.4536 (TOF
ESI+)

**[1]H NMR** ($CDCl_3$, 500 MHz)

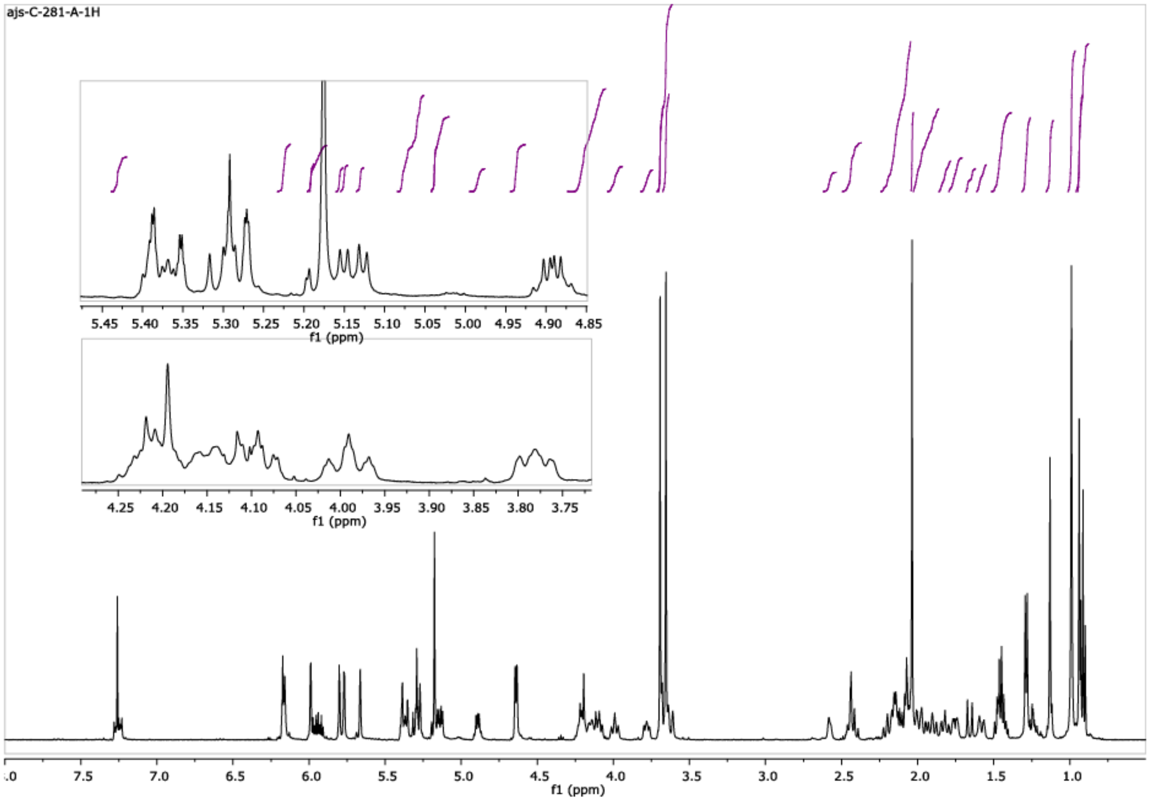


**[13]C NMR** ($CDCl_3$, 125 MHz)

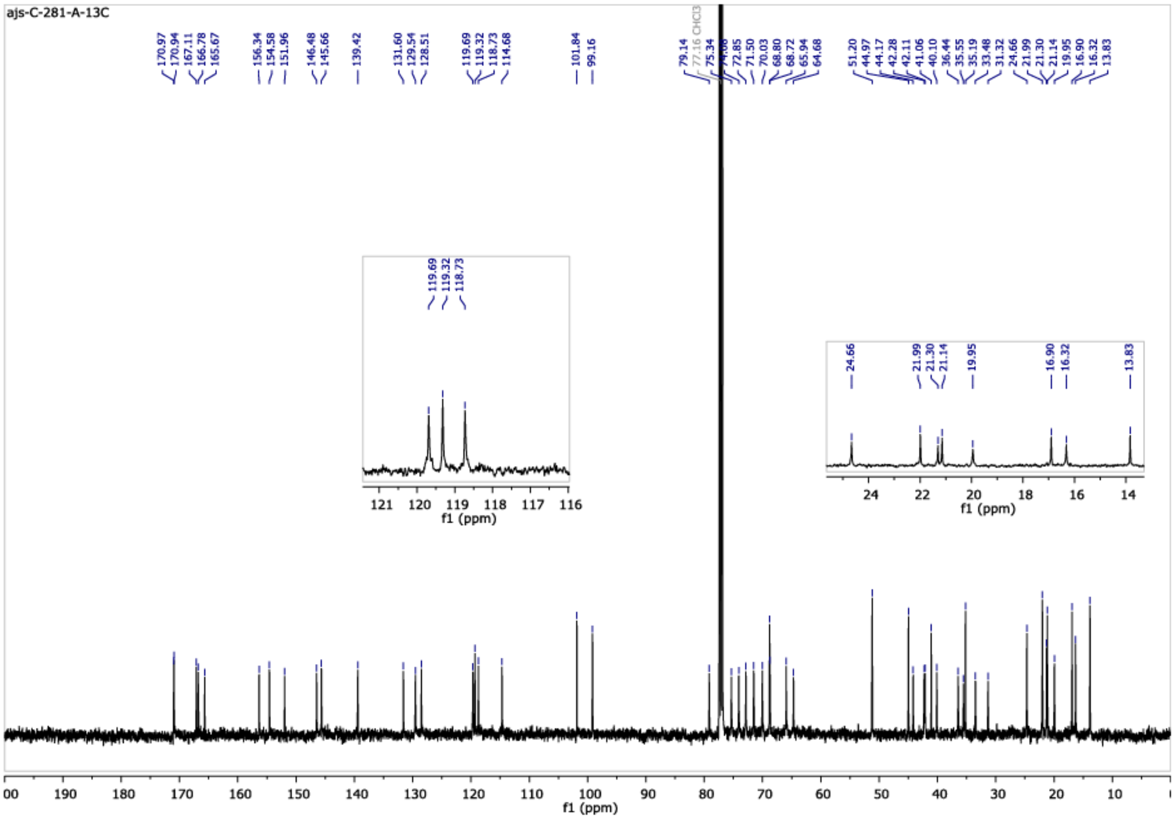

***Synthesis of C7-modified bryostatin analogs: Methanolysis of the C7 acetate***

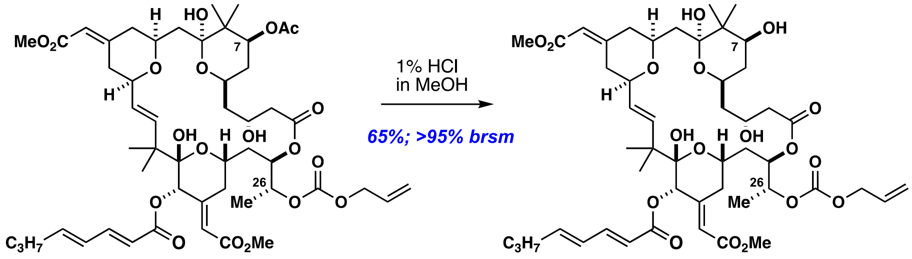


1

2

To a flame dried vial charged with a magnetic stir bar and 3.14 (6.2 mg, 0.0063 mmol) a solution of 1% HCl (vol/vol) in MeOH was added and the reaction was stirred at room temperature for 25 hours. The reaction solution was then quenched into 75 mL of water and extracted with DCM (5 portions of 75 mL). The combined organic layers were dried over Na2SO4, filtered, and concentrated to a crude film. Purification was accomplished by silica gel flash column chromatography (20%-40%-60%-100% EtOAc/Hex) affording product 3.15 (4.0 mg, 67% yield) and recovered starting material 3.14 (~2.0 mg, ~33% of starting material). Compound purity was established by TLC (one spot) analysis.

***Characterization data for C26-alloc protected C7 alcohol***

**TLC $R_f$** = 0.31 (60% EtOAc/hexanes, purple spot in p-anisaldehyde);

**$[\alpha]^{22.8}_D$** = 16.66° ± 0.55° (c = 0.10 wt/vol%, CH2Cl2);

IR (thin film) 3458, 3345, 2948, 2928, 2853, 1745, 1717, 1667, 1651, 1643, 1615, 1436, 1380, 1292, 1248, 1164, 1099, 1080, 1003, 859, 812 cm-1;

**$^1$H NMR** (600 MHz, Chloroform-d) δ 7.35 – 7.18 (m, 1H), 6.21 – 6.12 (m,2H), 6.00 (d, J = 2.0 Hz, 1H), 5.95 (ddt, J = 16.4, 10.9, 6.0 Hz, 1H), 5.80 (d, J = 2.5 Hz, 1H), 5.78 (d, J = 2.9 Hz, 1H), 5.67 (s, 1H), 5.44 – 5.26 (m, 4H), 5.19 (s, 1H), 5.17 (s, 1H), 4.90 (td, J = 6.8, 4.3 Hz, 1H), 4.64 (d, J = 4.5 Hz, 2H), 4.23 (d, J = 12.2 Hz, 1H), 4.17 – 4.06 (m, 3H), 4.04 – 3.94 (m, 2H), 3.81 – 3.75 (m, 1H), 3.70 (s, 3H), 3.66 (s, 3H), 3.63 (d, J = 14.4 Hz, 1H), 2.52 – 2.38 (m, 2H), 2.32 (s, 1H), 2.22 (t, J = 12.5 Hz, 1H), 2.16 (q, J = 6.7 Hz, 2H), 2.13 – 2.03 (m, 4H), 2.03 – 1.95 (m, 2H), 1.95 – 1.88 (m, 1H), 1.82 (t, J = 12.5 Hz, 1H), 1.72 – 1.56 (m, 4H), 1.48 – 1.42 (m, 3H), 1.29 (d, J = 6.4 Hz, 3H), 1.14 (s, 3H), 1.04 (s, 3H), 1.00 (s, 3H), 0.92 (t, J = 7.2 Hz, 3H), 0.91 – 0.89 (m, 3H);

**$^{13}$C NMR** (125 MHz, CDCl3) δ 171.1, 167.2, 166.8, 165.7, 156.5, 154.6, 152.0, 146.5, 145.7, 139.4, 131.6, 129.6, 128.5, 119.7, 119.4, 118.7, 114.7, 101.9, 99.2, 79.2, 75.4, 74.1, 71.6, 70.1, 70.0, 68.8, 68.7, 66.2, 64.7, 51.2, 51.2, 45.0, 44.3, 42.4, 42.4, 42.2, 40.2, 36.7, 36.5, 35.6, 35.2, 31.3, 24.7, 22.0, 21.2, 20.0, 16.4, 15.6, 13.9.;

**HRMS:** calculated for C49H70NaO18 [M+Na]+: 969.4454; found 969.4425 (TOF ESI+)

**[1]H NMR** ($CDCl_3$, 600 MHz)

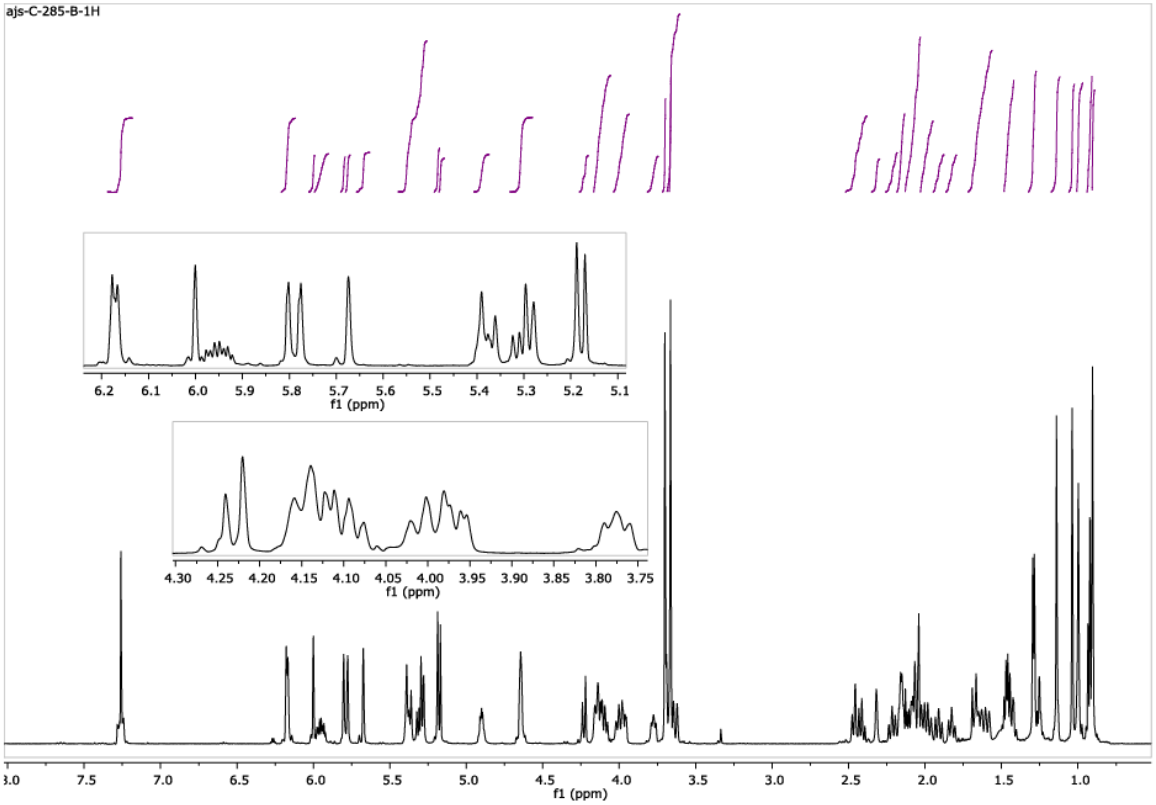


**[13]C NMR** ($CDCl_3$, 125 MHz)

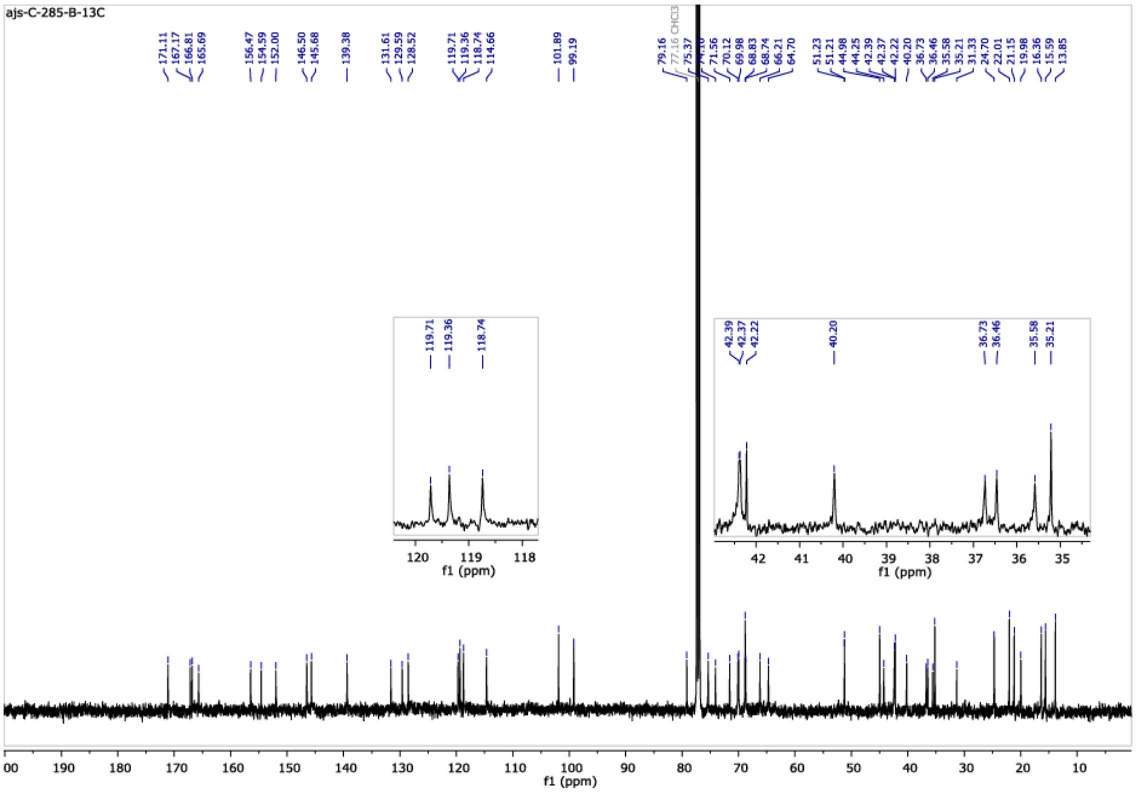

### *Synthesis of C7-modified bryostatin analog SUW239*

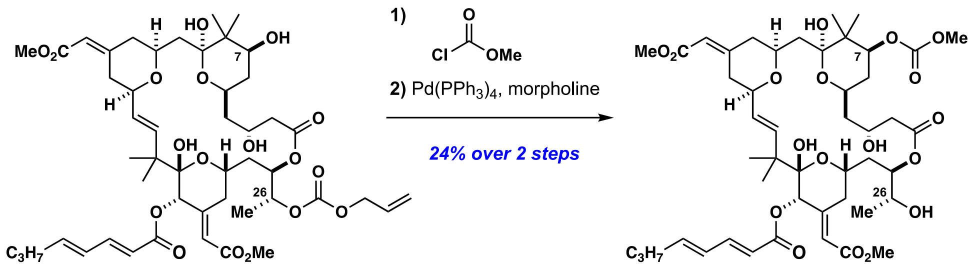


2 SUW239

*C7 carbonylation*

To a flame dried vial charged with a magnetic stir bar and 3.15 (2.5 mg, 0.0026 mmol) in DCM (0.2 mL), pyridine (2.1 µL, 0.026 mmol, ~10equiv.), methyl chloroformate (2 µL, 0.026 mmol, 10 equiv.) and DMAP (3.2 mg, 0.026 mmol, ~10 equiv.) were added in that order. The reaction was stirred for 24 hours, at which point TLC analysis showed little conversion. An additional portion of DMAP (3.2 mg, 0.026 mmol, ~10 equiv.) and methyl chloroformate (2 µL, 0.026 mmol, 10 equiv.) were added and the reaction was stirred for an additional 24 hours. The solution was then quenched into 10 mL of 0.1 N HCl and extracted with DCM (5 portions of 10 mL). The combined organic layers were dried over Na2SO4, filtered, and concentrated to a crude film. Purification was accomplished by silica gel flash column chromatography (20%-40%-60%-100% EtOAc/Hex) affording the carbonate (0.9 mg, 34% yield) and starting material 3.15 (1.6 mg, 64% of starting material). The intermediate product carbonate was obtained as a thin film and combined with product derived from the reaction being carried out a second time and then carried onto the next step.

*C26 Alloc deprotection*

The combined masses of the above reaction carried out twice (1.8 mg, 0.00179 mmol) was concentrated in a flame dried vial charged with a magnetic stir bar. To this thin film was added degassed (3 cycles of freeze-pump-thaw) THF (0.3 mL), morpholine (3.1 µL, 0.084 mmol, ~20 equiv.), and a solution of Pd(PPh3)4 in degassed THF (0.1 mL THF and 0.20 mg Pd(PPh3)4, 0.00042 mmol, ~0.1 equiv.). The resulting (yellow) solution was stirred for 24 hours. The solution was quenched into 10 mL of 0.02 N HCl and extracted with EtOAc (5 portions of 10 mL). The combined organic layers were dried over Na2SO4, filtered, and concentrated to a crude film. Purification was accomplished by silica gel flash column chromatography (40%-60%-100% EtOAc/Hex) affording the carbonate SUW239 (1.2 mg, 73% yield). The compound was additionally purified by semi-prep HPLC. Compound purity was established by TLC (one spot) analysis.

**Characterization data for SUW239**

**TLC $R_f$** = 0.32 (60% EtOAc/hexanes, purple spot in p-anisaldehyde);

**$[\alpha]^{24.1}_D$** = 37.6° ± 1.39° (c = 0.035 wt/vol%, CH2Cl2);

**IR** (thin film) 3465, 3345, 2955, 2924, 2851, 1716, 1699, 1668, 1661, 1652, 1436, 1272, 1248, 1164, 1080, 1097, 1058, 1004 cm-1;

**$^1$H NMR** (600 MHz, Chloroform-d) δ 7.28 – 7.20 (m, 1H), 6.17 – 6.12 (m, 2H), 5.98 (s, 1H), 5.78 (d, J = 15.2 Hz, 1H), 5.77 (d, J = 15.8 Hz, 1H), 5.66 (s, 1H), 5.29 (dd, J = 15.8, 8.3 Hz, 1H), 5.20 – 5.14 (m, 1H), 5.17 (s, 1H), 5.16 (s, 1H), 4.97 (dd, J = 11.8, 4.8 Hz, 1H), 4.25 – 4.11 (m, 3H), 4.04 (ddd, J = 11.1, 8.4, 2.2 Hz, 1H), 4.00 (tt, J = 11.2, 2.2 Hz, 1H), 3.82 – 3.75 (m, 2H), 3.74 (s, 3H), 3.68 (s, 3H), 3.65 (t, J = 13.3 Hz, 3H), 3.64 (s, 3H), 2.49 – 2.38 (m, 3H), 2.19 (t, J = 12.5 Hz, 1H), 2.16 – 2.11 (m, 2H), 2.11 – 1.77 (m, 9H), 1.65 (d, J = 15.1 Hz, 1H), 1.59 (dt, J = 15.0, 3.2 Hz, 1H), 1.44 (q,
J = 7.4 Hz, 2H), 1.22 (d, J = 6.4 Hz, 3H), 1.13 (s, 3H), 0.99 (s, 3H), 0.98 (s, 6H), 0.90 (t, J = 7.4 Hz, 3H).;

**$^{13}$C NMR** (125 MHz, CDCl3) δ 172.4, 167.2, 166.8, 165.7, 156.4, 155.7, 152.1, 146.5, 145.6, 139.4, 129.5, 128.5, 119.7, 118.8, 114.7, 102.0, 99.2, 79.3, 74.2, 73.9, 71.5, 70.3, 68.5, 66.0, 64.9, 64.8, 54.9, 51.2, 51.2, 45.1, 44.2, 42.5, 42.1, 41.4, 39.9, 36.4, 36.0, 35.2, 33.4, 31.4, 24.7, 22.0, 21.1, 20.0, 19.9, 16.8, 13.8.;

**HRMS:** calculated for C47H68NaO18 [M+Na]+: 943.4298; found 943.4297 (TOF ESI+)

**$^{1}$H NMR** (600 MHz, $CDCl_3$)

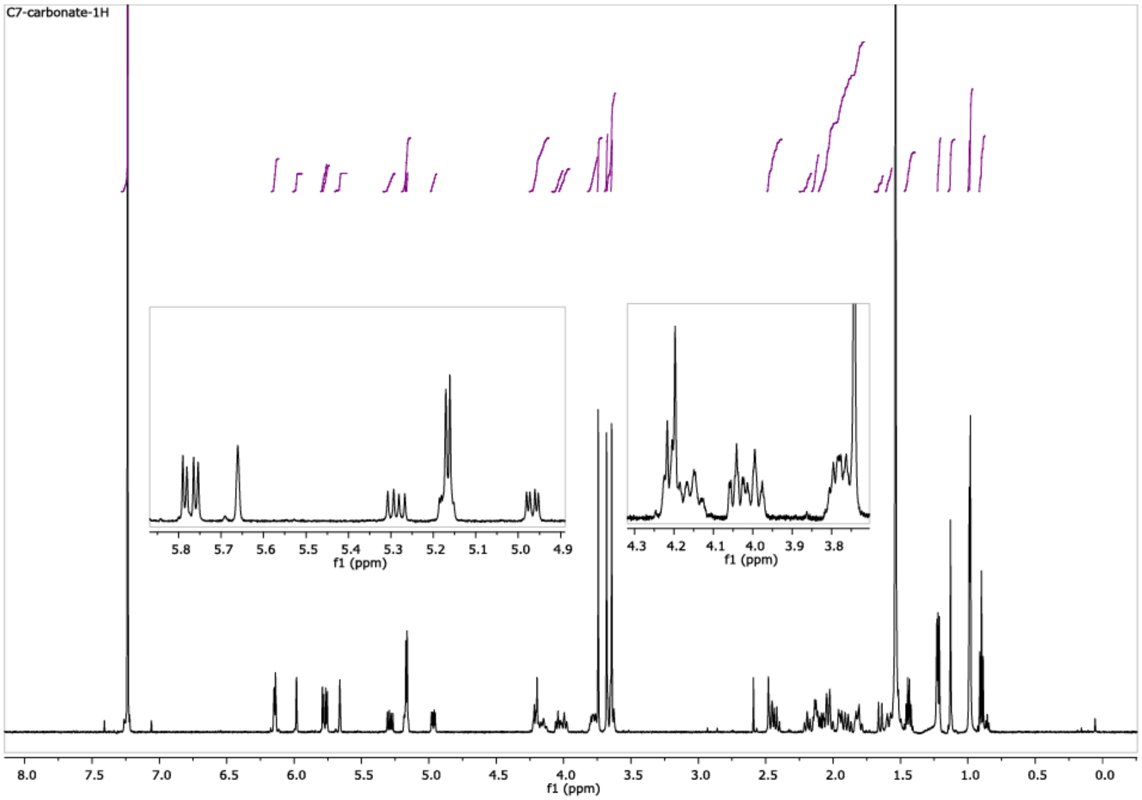


**$^{13}$C NMR** ($CDCl_3$, 125 MHz)

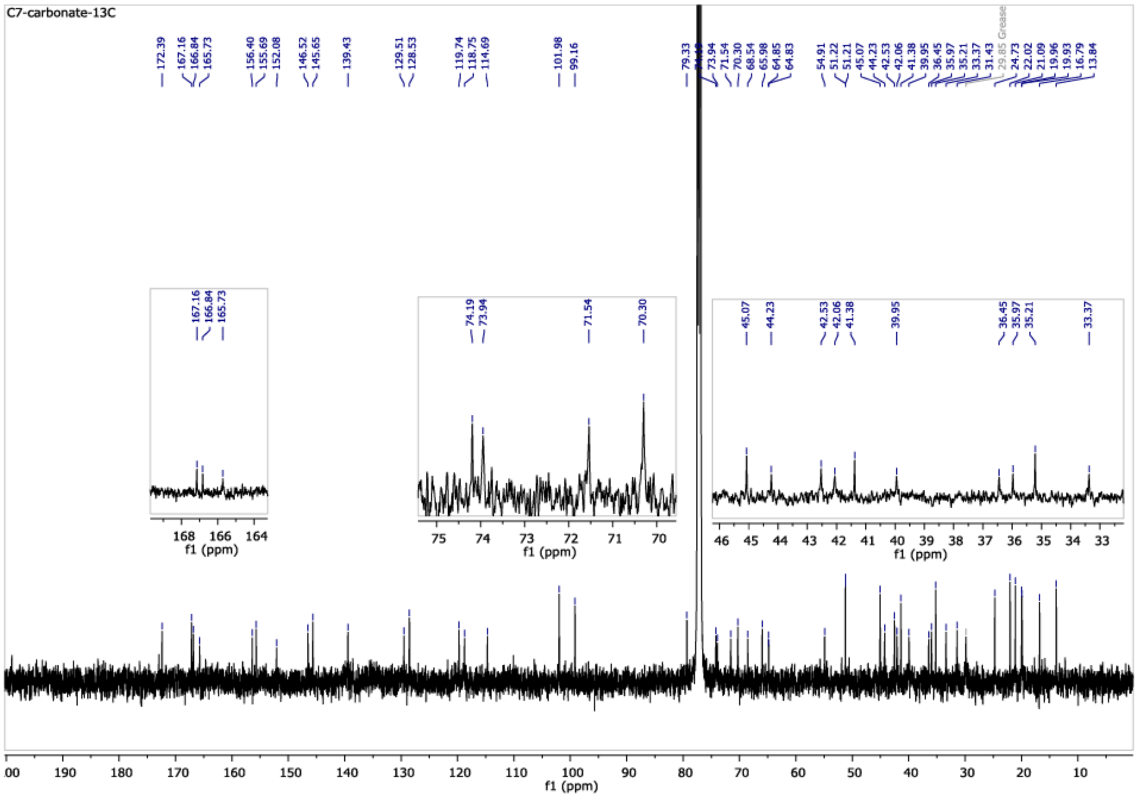
